# The ‘Threat of Scream’ paradigm: A tool for studying sustained physiological and subjective anxiety

**DOI:** 10.1101/834309

**Authors:** Morgan Beaurenaut, Elliot Tokarski, Guillaume Dezecache, Julie Grèzes

**Affiliations:** Laboratoire de Neurosciences Cognitives et Computationnelle, Département d’études cognitives, ENS, PSL Research University, INSERM, Paris France; Department of Experimental Psychology, Division of Psychology and Language Sciences, University College London, London, United Kingdom; Université Clermont Auvergne, CNRS, LAPSCO, Clermont-Ferrand, France

## Abstract

Progress in understanding the emergence of pathological anxiety depends on the availability of paradigms effective in inducing anxiety in a simple, consistent and sustained way. Much progress has been made using the Threat-of-Shock paradigm (TOS), which generates anxiety through the delivery of unpredictable electric shocks to participants. However, TOS may be problematic when testing vulnerable populations. Moreover, it is not clear whether anxiety can be sustained throughout experiments of long duration. Here, we bring support for an alternative approach called the Threat-of-Scream paradigm (TOSc), in which the tactile delivery of shocks is replaced by the auditory delivery of distress screams. We report on an one-hour long study (plus its replication) in which participants were exposed to blocks before which they were told that they could hear aversive screams at any time (threat blocks), vs. blocks before which they were told that no scream will be heard (safe blocks). Both the experiment and its replication showed higher subjective reports of anxiety, higher skin conductance level, and positive correlation between the two measures, in threat compared to safe blocks. Anxiety measures were sustained throughout the experiments, suggesting little emotional and physiological habituation. Our results suggest that the delivery of low intensity distress screams can be an efficient, stable and cheap methodology to assess the impact of sustained anxiety on a diversity of cognitive functions and populations. We therefore believe the TOSc will become an essential part of the psychological toolkit, particularly so for researchers interested in the emergence of pathological anxiety.

## Introduction

Given the ubiquity and persistence of anxiety disorders, as well as their massive impact upon quality of life (Leon, Portera, & Weissman, 1995), it is essential that neurobiologists and clinicians determine how anxiety influences human brain physiology and behavioural responses to stress or external threats, along the continuum from normal to pathological conditions. While high anxiety leads to exaggerated estimates of threat probability, a certain level of anxiety is crucial for an organism’s survival because it ensures optimal sensitivity and decisiveness in the face of possible threat (Bateson, Brilot, & Nettle, 2011; Grillon, 2008; Marks & Nesse, 1994)

The predictability of threats appears to be a major determinant of anxiety-related bodily manifestations, such as modulations of heart rate, startle reflex or skin conductance (Alvarez, Chen, Bodurka, Kaplan, & Grillon, 2011; Davis, Walker, Miles, & Grillon, 2010; Torrisi et al., 2016; Vansteenwegen, Iberico, Vervliet, Marescau, & Hermans, 2008). Indeed, predictable threats lead to phasic and acute fear responses that are directly associated with the appearance of a threat, e.g. a shock (e.g. startle reflex - Grillon, Baas, Lissek, Smith, & Milstein, 2004). In contrast, and in agreement with the safety-signal hypothesis (Seligman & Binik, 1977), unpredictable threats induce sustained anxiety-related physiological responses throughout the threat context (e.g. startle reflex - Grillon et al., 2004; prepulse inhibition, a physiological marker of alertness - Grillon & Davis, 1997) and enhanced vigilance, reflected by a long-lasting facilitated processing of sensory information (Kastner-Dorn, Andreatta, Pauli, & Wieser, 2018).

The Threat of Shock paradigm (hereafter TOS) has been the gold-standard paradigm to unveil the effects of anxiety on cognitive functions and quantify within-subject individual difference in threat response (for review - Robinson, Vytal, Cornwell, & Grillon, 2013). This paradigm consists in alternating blocks in which participants are explicitly told that they can receive electric shock at any time (unpredictable threat blocks) with blocks in which participants are explicitly told that no such shocks will occur (safe blocks). Such on-off alternations allow to manipulate the state of anxiety within subject as each participant can serve as her/his own control. The TOS paradigm proved to be a reliable method for inducing sustained anxiety, as reflected in participants’ physiological (startle reflex and elevated tonic skin conductance level) and psychological (higher reports of subjective anxiety) responses during threat versus safe blocks (Bradley, Zlatar, & Lang, 2017; Grillon, Robinson, Mathur, & Ernst, 2016; Hubbard et al., 2011; Torrisi et al., 2016)

Yet, and just like every paradigm, TOS has limitations. First, it is not clear whether it can induce sustained anxiety responses for long durations. Experiments that have employed the TOS paradigm have been relatively short (~ 30 minutes). One-hour long experiments using TOS do exist (Engelmann, Meyer, Fehr, & Ruff, 2015) but anxiety manipulation in those studies had mainly been assessed using self-rating of anxiety states, susceptible to demand effects, and/or using local phasic physiological changes, which only represent few seconds of the participants’ physiological state following the electric shock rather than sustained responses. To our knowledge, only two experiments looked at whether induced anxiety can be maintained across several blocks. Bublatzky and colleagues (Bublatzky, Guerra, Pastor, Schupp, & Vila, 2013) reported no habituation of tonic skin conductance activity but their experiment only lasted 15 minutes. Aylward and colleagues (Aylward et al., 2019) reported no habituation on subjective reports of anxiety throughout a 45-min experiment. It therefore remains unclear whether TOS is resistant to physiological habituation. Long-duration experiments are yet necessary when several conditions have to be tested and/or the analyses request a grand number of trials (as in computational modelling).

A second limitation has to do with the appropriateness of TOS for certain study populations. Although well-known for their aversive properties (Schmitz & Grillon, 2012), electric shocks may not be administrated to vulnerable and younger populations (notably children). The same critic arises for aversive noise, often presented between 95 and 110 dB, which is close to pain threshold and can cause hearing loss when prolonged (European Legislation-directive n° 2003/10/CE).

How to overcome such limitations, and best offer a methodology to generate anxiety in most of the populations, in a sustained and stable way? Threatening stimuli are useful insofar as they are perceived as unpleasant without being painful. As it stands, distress screams produced by humans are good candidates, as they are evolutionary and socially meaningful sounds that efficiently signal impending danger to conspecifics (Arnal, Flinker, Kleinschmidt, Giraud, & Poeppel, 2015; Belin & Zatorre, 2015). They are perceived as highly aversive signals and characterized by distinctive roughness acoustical properties (Anikin, Bååth, & Persson, 2018), which specifically engages subcortical regions known to be critical to swift reactions to danger (Arnal et al., 2015).

Human screams have previously been used during fear conditioning paradigms (e.g. screaming lady paradigm, J. Y. Lau et al., 2011; J. Y. F. Lau et al., 2008). Stimuli paired with screams induce higher skin conductance (Ahrens et al., 2016) and startle responses (Glenn, Lieberman, & Hajcak, 2012; Haddad, Xu, Raeder, & Lau, 2013) as well as higher subjective anxiety reports (Den, Graham, Newall, & Richardson, 2015) compared to unconditioned stimuli. As most of the past studies using screams delivered them with a potentially painful intensity (around or above 90 dB) (Ahrens et al., 2016; Dibbets & Evers, 2017; Geller et al., 2017; Hamm, Vaitl, & Lang, 1989; J. Y. Lau et al., 2011; J. Y. F. Lau et al., 2008), both the aversive nature of screams and the potential painful experience may contribute to the observed responses. Nevertheless, some studies succeeded to evoke acute fear response with intensity lower or equal to 80 dB (Den et al., 2015; Glenn, Klein, et al., 2012; Glenn, Lieberman, et al., 2012). Of interest, the fear potential startle was found to be comparable for stimuli conditioned with electric shocks and 80dB screams, even though those conditioned with shocks were reported to be more aversive than those conditioned with screams (Glenn, Lieberman, et al., 2012). If the substitution of shocks by screams is promising to generate acute stress, the question remains as to whether human screams can be an efficient tool to induce sustained anxiety in a within-subject paradigm. Of interest, Patel and colleagues (Patel, Stoodley, Pine, Grillon, & Ernst, 2017; Patel et al., 2016) manipulated anxiety using loud shrieking screams to be able to investigate working memory in adolescents (Threat of Scream paradigm – TOSc). Substituting shocks by screams was successful as participants reported being more anxious and had higher startle reflex in threat blocks compared to safe ones. Yet, the experiment only lasted thirty minutes and the screams were delivered at high intensity (95-dB).

To further establish the viability Threat of Scream paradigm (TOSc) and its promises for anxiety research, we used human distress screams delivered at low intensity -rather than high intensity screams or electric shocks- and tested their efficiency to induce anxiety during a one-hour experiment. Knowing that distress screams have specific acoustic properties and privileged communicative function, we expected them to be particularly suitable to evoke anxiety in a long-lasting fashion, when presented in an unpredictable manner. To determine whether sustained anxiety was induced, we measured subjective reports of anxiety and skin conductance activity, two markers that track environmental uncertainty (De Berker et al., 2016). We ran the same experiment of 1-hour duration twice, to assess whether the observed effects were reliable and replicable. Two conditions had to be met to validate the TOSc paradigm: (i) the unpredictable screams presented at low intensity (<80dB) should modulate anxiety responses, with increased subjective anxiety reports and increased tonic physiological activity (skin conductance) in threat compared to safe blocks, similarly to previous TOS studies; and (ii) sustained state anxiety should be induced for extended periods (here one hour), i.e. be resistant to habituation.

## Materials and Methods

### Participants

Twenty-six healthy volunteers (12 females, age 23.6 ± 3.4 years SD) were recruited to participate in Experiment 1 (sample size similar to Patel et al. (2016)’s study Threat of Scream). Results from the correlation between tonic skin conductance activity and subjective reports of anxiety were used to calculate the sample size needed to replicate this result using G*power. The sample size for replication was estimated at n = 27 for an effect size of d = 0.56, α = 0.05 and β = 0.80. To anticipate potential exclusions, we aimed at including 35 participants in Experiment 2 (Replication of Experiment 1), and 33 participants could be recruited (15 females, age = 23.89 ± 4.50).

All participants were right-handed, had normal or corrected-to-normal vision, and had no history of neurological or psychiatric disorders. The experimental protocol was approved by INSERM and the local research ethics committee (Comité de protection des personnes Ile de France III - Project CO7-28, N° Eudract: 207-A01125-48), and it was carried out in accordance with the Declaration of Helsinki. The participants provided informed written consent and were compensated for their participation.

### Questionnaires

Before the experiment, participants completed French online versions of STAI (State-Trait Anxiety Inventory Spielberger, Spielberger, 1983) and PCLS (Post-traumatic stress disorder Checklist Scale, Weathers, Litz, Herman, Huska, & Keane, 1993). Upon arrival at the laboratory, participants’ anxiety state was assessed once again using the STAI questionnaire. Due to the potentially stressful nature of our paradigm and after discussion with the referent medical doctors of our laboratory, we only recruited participants with a score below 60 for anxiety state and anxiety trait assessed by STAI and below 40 for PCLS.

### Procedure

The experiment was divided into 10 blocks, 5 threat blocks and 5 safe blocks, with an alternation between threat and safe blocks (Figure 1a). The nature of the first block was counterbalanced across participants. Participants were informed that when the sides of the screen were blue, they were at risk of hearing an unpredictable distress scream through their headphones (threat blocks); when the sides of the screen were green, no screams were to be delivered (safe blocks). During the 10 first seconds of each block, instructions relative to the nature of the block appeared on the screen, i.e. either “Threat Block: at any time, a scream can be presented” or “Safe Block: you will hear nothing during this block” (Figure 1b). During the 10 blocks, participants performed a free action-decision task in social context, developed by (Vilarem, Armony, & Grèzes, 2019), the data of this task being part of another study (Beaurenaut et al. unpublished).

**Figure 1:**
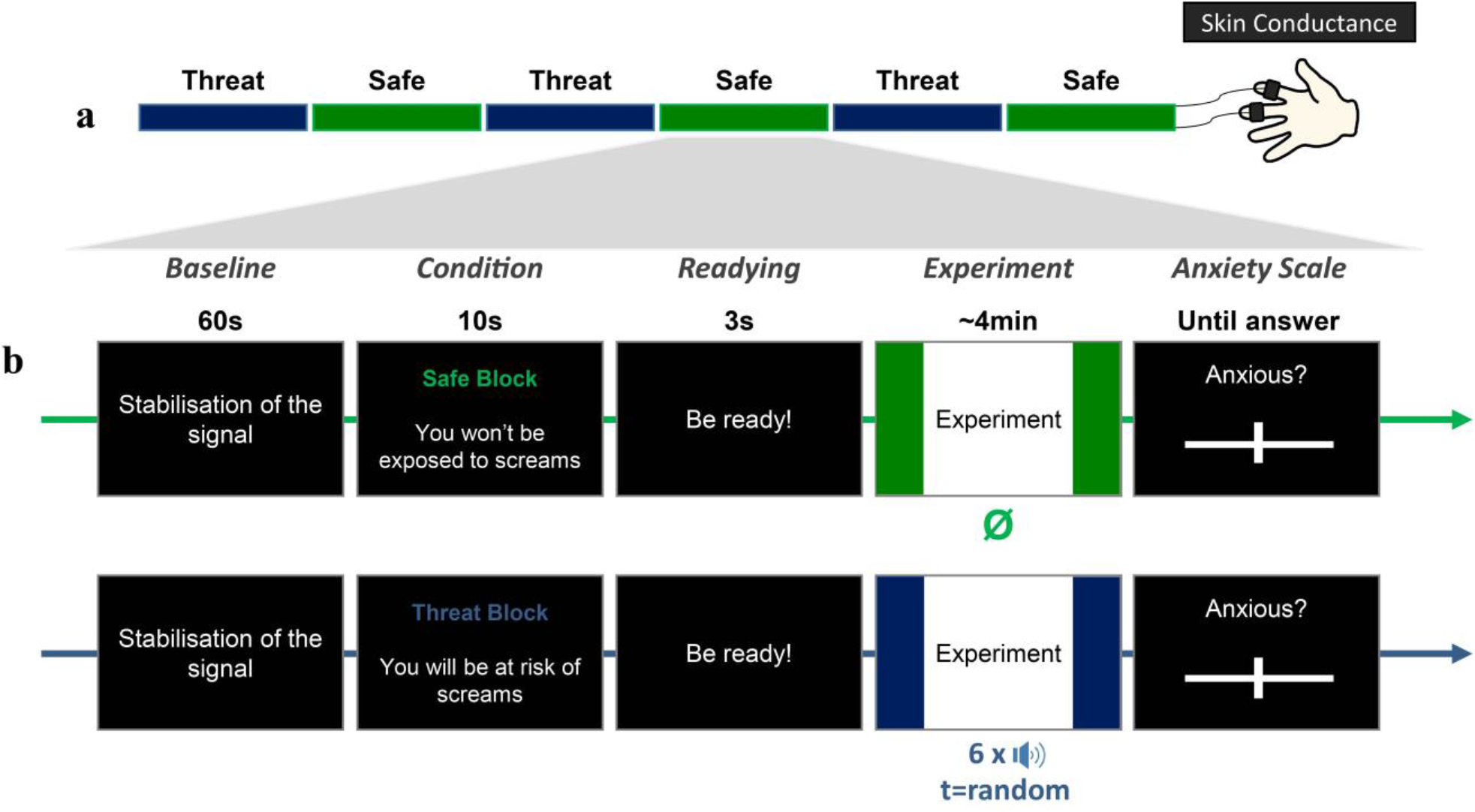
Illustration of the Threat of shock Design. (a) Temporal design of the experiment. Participants performed an experiment alternating the two types of blocks (Safe/Threat). (b) Temporal Design of a block. Each block began with one minute of baseline to assess the physiological state (skin conductance) of the participant before the experiment. Then the type of block was announced for 10s. Each block ended by an anxiety scale. The skin conductance activity was measured throughout the experiment whereas subjective anxiety ratings were collected at the end of each block.

To determine whether sustained anxiety was induced in threat as compared to safe blocks, skin conductance activity was recorded throughout the 1-hour experiment (M_duration_ = 65 min, SD_duration_ = 5min), whereas subjective reports of anxiety were retrospectively provided by participants at the end of each block, using a continuous scale (from 0: really calm to 100: really anxious).

### Training

A training session was performed before the main experiment to familiarize participants with the free action task, the structure of the experiment (safe and threat blocks depending on the colour of the screen sides) as well as with the type of screams that were to be delivered. The training was composed of 2 blocks, one safe block and one threat block, each composed of 32 trials.

### Screams stimuli

Eight distress screams were used in the present study (4 from males and 4 from females). They were provided by Professor Armony (Fecteau, Belin, Joanette, & Armony, 2007), and were normalized at −2b using audiosculpt 3.4.5 (http://forumnet.ircam.fr/shop/fr/forumnet/10-audiosculpt.html). During the experiment, the screams were delivered using Bose headphones (QuietComfort® 25) at peak intensity below 70 dB (mean of 68 dB as measured by a sonometer). During each experimental threat block, six distress screams were delivered once, randomly during the block (approximately 6% of the trials) either before the fixation, after the fixation or when the scene appeared. Similarly, during the training, 2 distress screams were delivered once. Note that the screams used for the training were different from the 6 screams used during the experimental phase.

### Skin conductance recordings

Physiological responses were recorded using ADInstruments© (ML870/Powerlab 8/30) acquisition system. Two Ag–AgCl electrodes filled with isotonic NaCl unbiase electrolyte were attached to the palmar surface of the middle phalanges of the index and middle fingers of the non-dominant hand. The skin conductance signals were recorded at a sampling rate of 1 kHz. Recordings were performed with Lab Chart 7 software, with the recording range set to 40 μS and using initial baseline correction (“subject zeroing”) to subtract the participant’s absolute level of electrodermal activity from all recordings. Finally, band-stop filter between 0.05 and 10Hz was applied on the signal.

### Skin conductance level (SCL) processing

The SCL corresponds to the tonic activity of the skin conductance signal. The processing of physiological signal was performed using Labchart 7 and Matlab. The SCL baseline was calculated using the minute break before the beginning of each block (see Figure 1B). The signal was averaged across each 4 minutes block (M_Duration_ = 4min, SD = 30s), deviated from this 1-min baseline. The phasic activity induced by the distress scream where excluded from the averaged signal, corresponding to around 6% of threat trials. Variations from baseline were obtained, corresponding to modifications of the Skin conductance activity after the start of the anxiety manipulation (either a threat or safe block). Variations from baseline were z-scored within-subject to account for individual differences.

### Statistical analyses

All statistical tests were carried out using JASP Software (JASP Team (2017), JASP (Version 0.8.5.1) [Computer software]). All the Tabs were available in the supplementary materials.

#### Repeated-measures ANOVAs

For each experiment and for physiological variables (SCL) and subjective reports of anxiety, we ran one-way repeated-measures ANOVA with Condition (Threat vs. Safe) and Time (block 1 to 5) as within-subject factor. We applied the Greenhouse-Geisser correction to correct for deviations from the assumption of sphericity (the corrected *P corrected* are reported) and Bonferroni correction for multiple comparisons (P*bonf*). Effect sizes (eta-squared, η^2^) are reported together with F and p values. As we expected, from previous findings using the Threat-of-Shocks paradigm, higher levels of physiological responses and of subjective reports of anxiety in Threat compared to Safe contexts, we conducted unilateral t-tests on significant main effects. Effect sizes (Cohen’s d) are reported together with t and p values.

#### Intra-individual correlation

To assess the intra-individual coherence between the physiological state of participants and their subjective experience, we first computed Pearson‘s r correlation coefficient for each participant between their subjective reports and SCL measures (10 values for each measure and each participant, since the experiment was composed of 10 blocks). We then performed Fisher’s r‐to‐z transformation to normalize Pearson’s r correlation coefficients (Howell, 2012) before testing whether the correlation coefficients across participants were different from zero (bidirectional one-sample t-test).

## Results Experiment 1

### Skin conductance level (SCL)

Participants’ tonic skin conductance activity (SCL) was greater during Threat relative to Safe blocks (F(1,25) = 23.81, p < 0.001, ƞ^2^ = 0.49) (**Figure 2a**). There was no significant interaction between Time and Condition (F(4,100) = 1.78, p = 0.14, p_corr_ = 0.16, ƞ^2^ = 0.067). Unilateral t-test and Cohen’s d effect size showed that the significant difference in SCL between Threat and Safe blocks was large (t(25) = 4.88, p < 0.001, Cohen’s d = 0.96 and lower 95% CI for Cohen’s d = 0.56).

**Figure 2:**
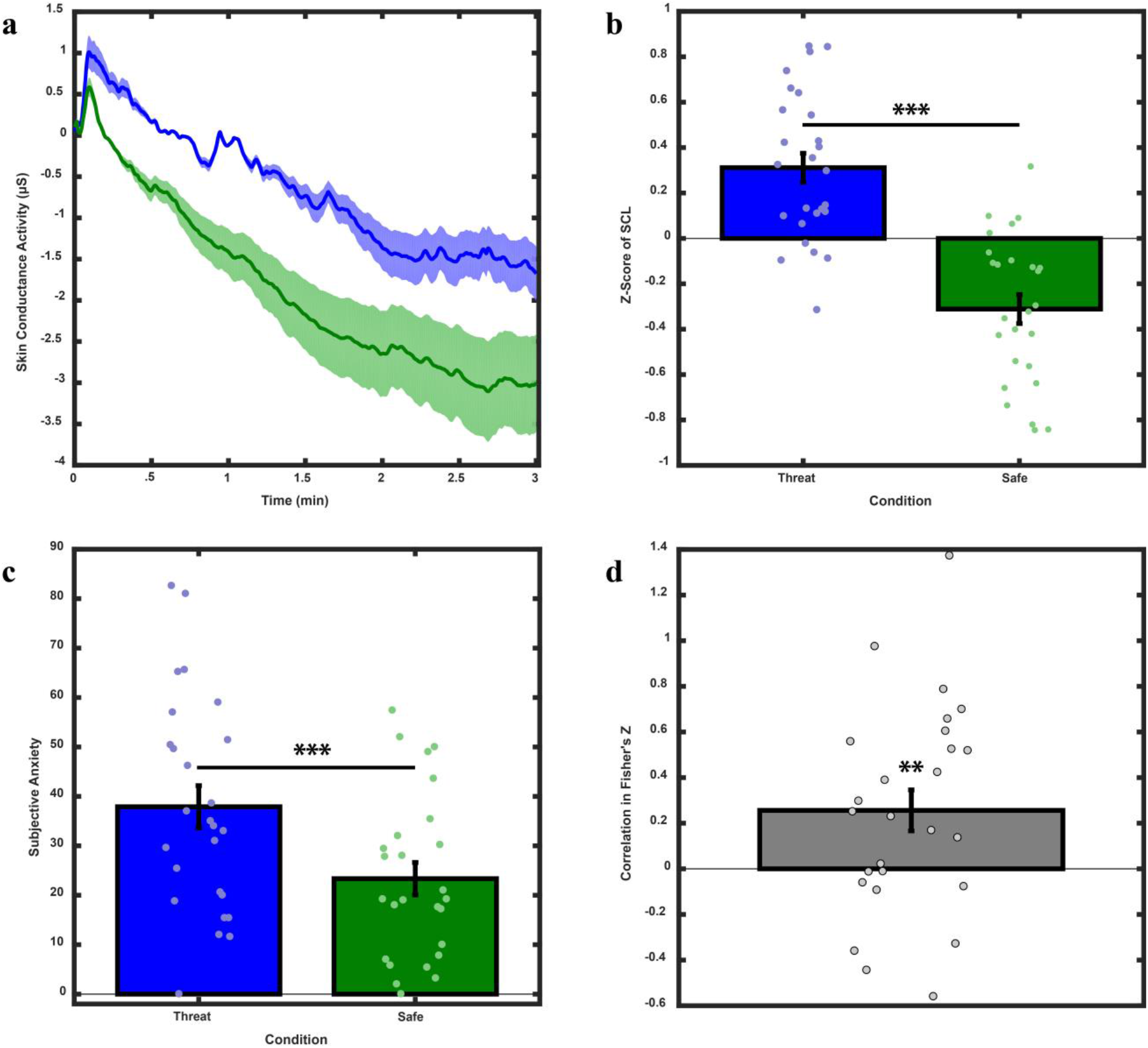
Results from Experiment 1. (**a**) Grand mean of skin conductance activity across the duration of one block, for threat and safe conditions separately. Shaded error bars indicate SEM. (b) Z-score of the mean SCL during Threat and Safe blocks (+/− SEM). Mean Difference of Z-Score SCL = 0.62, SE = 0.13. (**c**) Mean of subjective reports of anxiety during Threat and Safe blocks (+/− SEM). Mean Threat = 37.9, SE = 4.27; Mean Safe = 23.35, SE = 3.28. (**d**) Mean intra-individual correlation between SCL and subjective reports of anxiety. Mean = 0.26, SE = 0.09. *** = p < 0.001; ** = p < 0.01; * = p < 0.05; n.s. = p > 0.05.

### Subjective reports of anxiety

Participants reported higher scores on the anxiety scale at the end of Threat blocks compared to the end of Safe ones (F(1,25) = 15.11, p < 0.001, ƞ^2^ = 0.38) (**Figure 2b**). There is a trend toward significance for the interaction between Time and Condition on subjective reports (F(4,100) = 2.14, p = 0.081, ƞ^2^ = 0.079). Unilateral t-test and Cohen’s d effect size showed that the difference in subjective anxiety between Threat and Safe blocks was medium (t(25) = 3.89, p < 0.001, Cohen’s d = 0.76 and lower 95% CI for Cohen’s d = 0.39).

### Intra-individual Correlation

The average of intra-individual correlation estimates between SCL and subjective reports was positive, of medium size and statistically different from zero (bidirectional t-test, t(25) = 2.86, p = 0.009, Cohen’s d = 0.56, lower 95% CI for Cohen’s d = 0.14, upper 95% CI for Cohen’s d = 0.97) (**Figure 2c**).

## Results Experiment 2 (Replication)

### Skin conductance level (SCL)

Replicating results from Experiment 1, participants’ tonic skin conductance activity (SCL) was higher during Threat relative to Safe blocks (F(1,32) = 35.31, p < 0.001, ƞ^2^ = 0.53). There was no significant interaction between Time and Condition (F(4,128) = 1.49, p = 0.21, ƞ^2^ = 0.044). Unilateral t-test and Cohen’s d effect size showed that the difference in SCL between Threat and Safe blocks was large (t(32) = 5.94, p < 0.001, Cohen’s d = 1.03 and lower 95% CI for Cohen’s d = 0.67) (**Figure 3a**).

**Figure 3:**
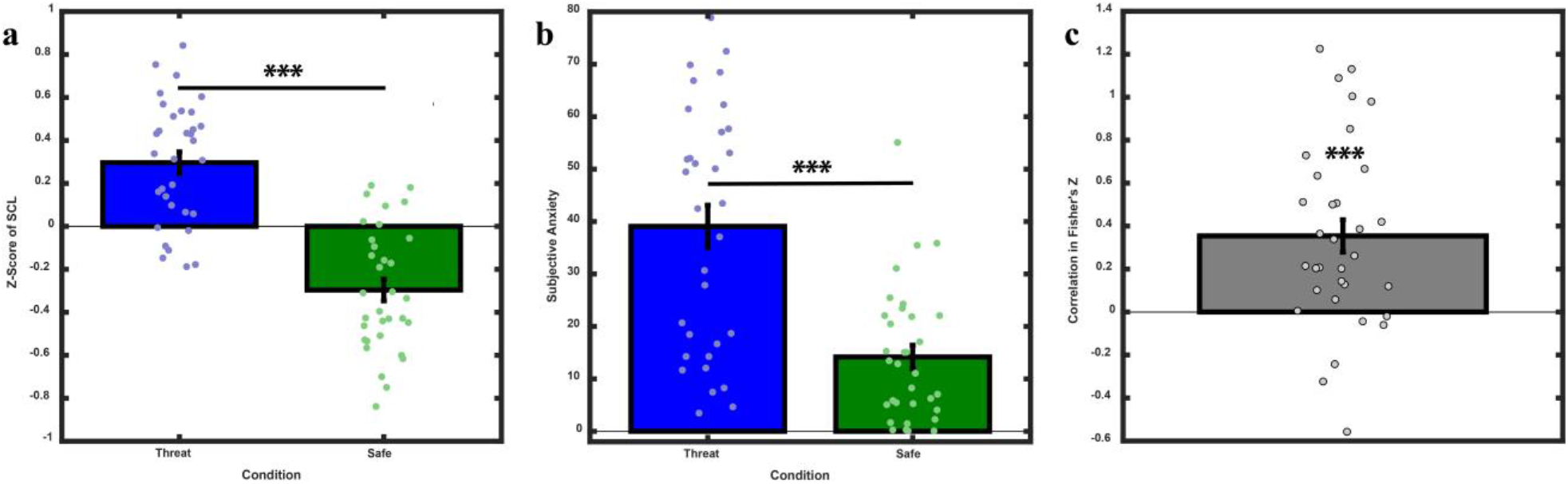
Results from Experiment 2. (**a**) Z-score of the mean SCL during Threat and Safe blocks (+/− SEM). Mean Difference of Z-Score SCL = 0.59, SE = 0.10. (**b**) Mean of Subjective reports of anxiety during Threat and Safe blocks (+/− SEM). Mean Threat = 39.08, SE = 4.04; Mean Safe = 14.19, SE = 2.23. (**c**) Mean of intra-individual correlation between SCL and Subjective reports of anxiety. Mean = 0.35, SE = 0.075. *** = p < 0.001; ** = p < 0.01; * = p < 0.05; n.s. = p > 0.05.

### Subjective reports of anxiety

Replicating results from Experiment 1, participants reported higher scores on the anxiety scale at the end of Threat compared to Safe blocks (F(1,32) = 47.84, p < 0.001, ƞ^2^ = 0.60) (**Figure 3b**). There was a significant interaction between Condition and Time on subjective reports (F(4,128) = 4.046, p = 0.004, ƞ^2^ = 0.112). A post-hoc test revealed a significant decrease in subjective reports between the first and the last threat blocks (Mean Difference= 13.27, SE = 3.89, t(32) = 3.41, p_bonf_ = 0.018). However, the contrast between each Safe and Threat block was significant (all the p_bonf_ < 0.001, 0.77 < Cohen’s d < 1.24) (See **Supplementary Figure S1**). Unilateral t-test and Cohen’s d effect size showed that the difference in subjective anxiety between Threat and Safe blocks was large (t(32) = 6.92, p < 0.001, Cohen’s d = 1.20 and lower 95% CI for Cohen’s d = 0.82).

### Intra-individual Correlation

Replicating results from Experiment 1, mean of intra-individual correlation between SCL and subjective reports of anxiety was positive, of large size and statistically different from zero (Bidirectional T-test-t(32) = 4.70, p < 0.001, Cohen’s d = 0.82, lower 95% CI for Cohen’s d = 0.42 upper 95% CI for Cohen’s d = 1.21) (**Figure 3c**).

## Discussion

The present experiment aimed at further validating an existing Threat-of-Scream paradigm (TOSc - Patel et al., 2016) by investigating whether sustained anxiety can be induced without habituation during 1 hour, using unpredictable distress screams delivered at low sensory intensity (70-dB). We measured two proxies of anxiety, namely subjective reports of anxiety and skin conductance level, and ran the same 1-hour duration experiment twice to assess the reliability and replicability of our results. Both experiments revealed higher skin conductance level (SCL) and self-reported anxiety during threat blocks compared to safe ones. Moreover, we observed no habituation effect on SCL, though participant reported less anxiety at the end of the experiment. Physiological state of participants (SCL) and their subjective reports of anxiety were positively correlated and all reported effect sizes were medium to large and replicable. Overall, our findings bring convincing evidence that the TOSc paradigm is a robust tool to assess the relatively long-lasting impact of sustained anxiety, within subject, with potential applications to a diversity of cognitive functions and populations.

### Human distress screams efficiently induce sustained anxiety on prolonged period of time

To manipulate anxiety within participants, we used unpredictable human distress screams as aversive cues. Human distress screams are highly salient vocal signals of impending danger. Characterized by distinctive roughness acoustical properties which contribute to their aversiveness (Arnal et al., 2015; Arnal, Kleinschmidt, Spinelli, Giraud, & Mégevand, 2019), they are perceived as communicating fear (Anikin, Bååth, & Persson, 2018) and convey cues to caller identity (Engelberg, Schwartz, & Gouzoules, 2019). Past experiments successfully substituted shocks by human screams to generate acute stress during fear conditioning paradigms (e.g. screaming lady paradigm, J. Y. Lau et al., 2011; J. Y. F. Lau et al., 2008). For instance, comparable physiological responses (fear potential startle) were found for stimuli conditioned with electric shocks and 80dB screams, even if stimuli conditioned with electric shocks were rated as more aversive than those conditioned with screams (Glenn, Lieberman, et al., 2012).

Knowing that unpredictability can induce sustained anxiety providing that the anticipated stimulus is sufficiently aversive (Grillon et al., 2004), we tested here whether unpredictable distress screams, delivered at lower sensory intensity (70-dB) than previous experiments (Patel et al., 2017, 2016), can be an efficient tool to induce sustained anxiety. Our findings clearly demonstrate that the aversiveness of unpredictable distress screams at low sensory intensity (70-dB) is enough to induce sustained anxiety in a within-subject paradigm. Indeed, in addition to a modest but significant increase in self-reported anxiety, participants’ skin conductance level increased in blocks during which human distress screams were delivered compared to safe blocks, and this difference in physiological activity between threat and safe blocks was large in both experiments (Cohen’s d>0.95).

Moreover, we demonstrate that anxiety can be induced for extended periods (here one hour) and appears resistant to habituation. Indeed, we observed in two experiments that changes between threat and safe blocks in both self-reported anxiety and tonic physiological activity (skin conductance level) remained constant over time. Our results extend previous findings of sustained tonic skin conductance activity during 15-min study (Bublatzky et al., 2013) and of self-reported anxiety throughout a 45-min experiment (Aylward et al., 2019). In agreement with the safety-signal hypothesis (Seligman & Binik 1977), the on-off alternation between threat and safe contexts evoked an on-off heightened state of vigilance in participants, reflected by increased self-reported anxiety and skin conductance level, a phenomenon which can be maintained with no habituation for extended periods of time. Altogether, findings from the current study indicate that unpredictable distress screams, which serve to communicate danger, i) are efficient to manipulate anxiety within-subject for at least one hour, and ii) they can be perceived as aversive even at low intensity.

### Correlation between subjective and physiological responses

Subjective reports of anxiety and skin conductance level were found to be positively correlated in both experiments. The association between self-reported experience and observed physiological activity was labelled in the past “emotion coherence”, and many theorists have suggested that coordinated changes (coherence) across physiological, behavioural (facial expressions), and experiential responses is the definition of an emotion episode. This proposition is however debated, as some authors suggested that emotion systems are only loosely coupled (Bonanno & Keltner, 2004; Izard, 1977), and, more recently, could even be independent (e.g., Ledoux, Ph, & Pine, 2016). Results from past experiments in healthy subjects are inconclusive as they provided evidence either for a moderate association between physiological responses and self-rated experience (Cuthbert, Schupp, Bradley, Birbaumer, & Lang, 2000; Dan-Glauser & Gross, 2013; Franklin et al., 2016; Mauss, Levenson, McCarter, Wilhelm, & Gross, 2005; Nandrino et al., 2012) or were consistent with the hypothesis that there is no emotional coherence between subjective and physiological data (Morris, DeGelder, Weiskrantz, & Dolan, 2001; Vuilleumier, Armony, Driver, & Dolan, 2001; Vuilleumier & Pourtois, 2007).

Within the fear conditioning literature, positive correlations between SCR and self-reported experience have been observed (Lovibond, Davis, & O’Flaherty, 2000; Rodriguez, Craske, Mineka, & Hladek, 1999). For instance, Glenn, Lieberman, Hajcak (2012) observed at a trend level a convergence between subjective and physiological (fear-potentiated startle) measures of fear, but only for stimuli that were conditioned using electric shocks and not for those conditioned with screams. By contrast, by alternating blocks during which participants were at risk of hearing unpredictable aversive screams with blocks during which no screams were to be delivered (safe blocks), we show that unpredictable distress screams at low sensory intensity (70-dB) are sufficiently aversive to generate emotion coherence, i.e. positive correlation between subjective reports of anxiety and skin conductance level. Such coordinated changes (coherence) across physiological and experiential responses clearly support the efficiency of our manipulation to induce an emotion (anxiety) episode.

## Conclusion

Overall, our study offers support to the TOSc paradigm by showing that: (i) unpredictable distress screams presented at low intensity (<80dB) can induce sustained anxiety as revealed by increased subjective reports of anxiety and increased tonic physiological activity (skin conductance) during threat compared to safe blocks, similarly to previous Threat of Shock experiments; and that (ii) sustained state anxiety can be induced for extended periods (here one hour), i.e. with some resistance to habituation. Distress screams, delivered at lower sensory intensity (70-dB), thus appear to be excellent candidates to overcome the ethical issues associated with exposing vulnerable and young populations to electric shocks and aversive noise, and to experimentally address the emergence of pathological anxiety in a consistent and sustained manner.

## Author notes

### Author contributions

MB, JG and GD developed the Threat of Scream paradigm. MB conducted the experiments and performed data analysis with the help of ET. MB, JG and GD wrote the first draft of the manuscript. All authors reviewed and approved the final manuscript.

## Acknowledgments

This work was supported by FRM Team DEQ20160334878, Fondation ROGER DE SPOELBERCH, INSERM, ENS, the French National Research Agency under Grants ANR-10-LABX-0087 IEC, ANR-10-IDEX-0001-02 PSL*, and ANR-17-EURE-0017 FrontCog. Author GD is indebted to the British Academy for financial support as part of the Newton International fellowship scheme. We thank Jorge Armory for sharing his distress screams and the *<u>Perception et design sonores</u>* team (working at *Institut de Recherche et Coordination Acoustique/Musique, Paris, France)* for their assistance.

## Competing interests

The authors declare no competing financial interests.

