## Supplementary materials for "The ‘Threat of Scream’ paradigm: A tool for studying sustained physiological and subjective anxiety"

### Description

The present work aimed at assessing the efficiency of a new version of TOSc paradigm during which we delivered unpredictable human distress screams at low intensity (70dB instead of 95dB used in the past). In the main manuscript, we discussed findings related to tonic skin conductance activity (which represent the physiological activity of participant through the entire block) and subjective scores of anxiety reported by participants at the end of each block. This supplementary document provides detailed results for all the analyses reported in the main text (Descriptive Statistics, Repeated Measures ANOVA, Unilateral Paired Samples T-Test, Post Hoc Comparisons and One Sample T-Test Against Zero)

### Experiment 1 – Supplementary results

***Skin Conductance Level***

**Table S1.** Within Subjects Effects of Repeated Measures ANOVA (Condition & Time) on SCL

| | Sphericity Correction | Sum of Squares | df | Mean Square | F | p | η² |
|---|---|---|---|---|---|---|---|
| Condition | None | 25.164 | 1.000 | 25.164 | 23.809 | < .001 | 0.488 |
| | Greenhouse-Geisser | 25.164 | 1.000 | 25.164 | 23.809 | < .001 | 0.488 |
| Residual | None | 26.423 | 25.000 | 1.057 | | | |
| | Greenhouse-Geisser | 26.423 | 25.000 | 1.057 | | | |
| Time | None | 44.703 | 4.000 | 11.176 | 18.365 | < .001 | 0.423 |
| | Greenhouse-Geisser | 44.703 | 3.401 | 13.144 | 18.365 | < .001 | 0.423 |
| Residual | None | 60.854 | 100.000 | 0.609 | | | |
| | Greenhouse-Geisser | 60.854 | 85.025 | 0.716 | | | |
| Condition ✻ Time | None | 5.118 [a] | 4.000 [a] | 1.279 [a] | 1.784 [a] | 0.138 [a] | 0.067 |
| | Greenhouse-Geisser | 5.118 [a] | 2.866 [a] | 1.786 [a] | 1.784 [a] | 0.160 [a] | 0.067 |
| Residual | None | 71.737 | 100.000 | 0.717 | | | |
| | Greenhouse-Geisser | 71.737 | 71.658 | 1.001 | | | |

*Note.* Type III Sum of Squares

[a] Mauchly's test of sphericity indicates that the assumption of sphericity is violated (p < .05).

**Table S2.** Descriptive Statistics (ANOVA) - SCL

| Condition | Time | Mean | SD | N |
|---|---|---|---|---|
| Threat | 1 | 1.297 | 0.805 | 26 |
| | 2 | 0.304 | 0.711 | 26 |
| | 3 | 0.296 | 0.787 | 26 |
| | 4 | 0.010 | 0.710 | 26 |
| | 5 | -0.352 | 0.749 | 26 |
| Safe | 1 | 0.255 | 1.275 | 26 |
| | 2 | -0.395 | 0.529 | 26 |
| | 3 | -0.413 | 0.763 | 26 |
| | 4 | -0.451 | 0.670 | 26 |
| | 5 | -0.552 | 0.765 | 26 |

**Table S3.** Unilateral Paired Samples T-Test *(Condition)* for SCL

| Condition | | t | df | p | Mean Difference | SE Difference | 95% CI for Mean Difference | | Cohen's d | 95% CI for Cohen's d | |
|---|---|---|---|---|---|---|---|---|---|---|---|
| | | | | | | | Lower | Upper | | Lower | Upper |
| Threat | - Safe | 4.879 | 25 | < .001 | 0.622 | 0.128 | 0.404 | ∞ | 0.957 | 0.558 | ∞ |

*Note.* Student's t-test.

*Note.* All tests, hypothesis is measurement one greater than measurement two.

**Table S4.** Descriptive Statistics (T-Test) - SCL

| Condition | N | Mean | SD | SE |
|---|---|---|---|---|
| Threat | 26 | 0.311 | 0.325 | 0.064 |
| Safe | 26 | -0.311 | 0.325 | 0.064 |

***Subjective reports of Anxiety***

**Table S5.** Within Subjects Effects of Repeated Measures ANOVA *(Condition & Time)* on Subjective Anxiety

| | Sphericity Correction | Sum of Squares | df | Mean Square | F | p | $\eta^2$ |
|---|---|---|---|---|---|---|---|
| Condition | None | 13753.4 | 1.000 | 13753.39 | 15.107 | < .001 | 0.377 |
| | Greenhouse-Geisser | 13753.4 | 1.000 | 13753.39 | 15.107 | < .001 | 0.377 |
| Residual | None | 22760.3 | 25.000 | 910.41 | | | |
| | Greenhouse-Geisser | 22760.3 | 25.000 | 910.41 | | | |
| Time | None | 1046.9 [a] | 4.000 [a] | 261.74 [a] | 1.841 [a] | 0.127 [a] | 0.069 |
| | Greenhouse-Geisser | 1046.9 [a] | 2.434 [a] | 430.05 [a] | 1.841 [a] | 0.159 [a] | 0.069 |
| Residual | None | 14217.7 | 100.000 | 142.18 | | | |
| | Greenhouse-Geisser | 14217.7 | 60.862 | 233.61 | | | |
| Condition ✻ Time | None | 793.0 | 4.000 | 198.24 | 2.143 | 0.081 | 0.079 |
| | Greenhouse-Geisser | 793.0 | 2.883 | 275.05 | 2.143 | 0.105 | 0.079 |
| Residual | None | 9252.8 | 100.000 | 92.53 | | | |
| | Greenhouse-Geisser | 9252.8 | 72.075 | 128.38 | | | |

*Note.* Type III Sum of Squares

[a] Mauchly's test of sphericity indicates that the assumption of sphericity is violated ($p < .05$).

**Table S6.** Descriptive Statistics (ANOVA) - Subjective Anxiety

| Condition | Time | Mean | SD | N |
|---|---|---|---|---|
| Threat | 1 | 43.00 | 23.09 | 26 |
| | 2 | 39.12 | 24.44 | 26 |
| | 3 | 37.69 | 23.31 | 26 |
| | 4 | 36.73 | 25.37 | 26 |
| | 5 | 32.96 | 25.68 | 26 |
| Safe | 1 | 22.35 | 19.78 | 26 |
| | 2 | 25.81 | 20.15 | 26 |
| | 3 | 25.27 | 17.68 | 26 |
| | 4 | 20.92 | 17.20 | 26 |
| | 5 | 22.42 | 17.94 | 26 |

**Table S7.** Unilateral Paired Samples T-Test (Condition) for Subjective Anxiety

| | | | | | | | 95% CI for Mean Difference | | | 95% CI for Cohen's d | |
|---|---|---|---|---|---|---|---|---|---|---|---|
| Condition | | t | df | p | Mean Difference | SE Difference | Lower | Upper | Cohen's d | Lower | Upper |
| Threat | - Safe | 3.887 | 25 | < .001 | 14.55 | 3.743 | 8.153 | ∞ | 0.762 | 0.388 | ∞ |

*Note.* Student's t-test.

*Note.* All tests, hypothesis is measurement one greater than measurement two.

**Table S8.** Descriptive Statistics (T-Test) - Subjective Anxiety

| Condition | N | Mean | SD | SE |
|---|---|---|---|---|
| Threat | 26 | 37.90 | 21.76 | 4.268 |
| Safe | 26 | 23.35 | 16.72 | 3.278 |

***Intra-individual Correlation***

**Table S9.** One Sample T-Test for Intra-individual Correlation (r to z Fischer)

| | | | | | 95% CI for Mean Difference | | | 95% CI for Cohen's d | |
|---|---|---|---|---|---|---|---|---|---|
| | t | df | p | Mean Difference | Lower | Upper | Cohen's d | Lower | Upper |
| Correlation | 2.855 | 25 | 0.009 | 0.256 | 0.071 | 0.441 | 0.560 | 0.141 | 0.969 |

*Note.* Student's t-test.

**Table S10.** Descriptive Statistics (T-Test) – Intra-individual Correlation

| | N | Mean | SD | SE |
|---|---|---|---|---|
| Correlation | 26.00 | 0.256 | 0.458 | 0.090 |

### Experiment 2– Supplementary results

##### *Skin conductance level (SCL)*

**Table S11.** Within Subjects Effects of Repeated Measures ANOVA *(Condition & Time)* on SCL

| | Sum of Squares | df | Mean Square | F | p | η² |
|---|---|---|---|---|---|---|
| Condition | 28.974 | 1 | 28.974 | 35.305 | < .001 | 0.525 |
| Residual | 26.262 | 32 | 0.821 | | | |
| Time | 28.612 | 4 | 7.153 | 8.026 | < .001 | 0.201 |
| Residual | 114.073 | 128 | 0.891 | | | |
| Condition ✻ Time | 4.397 | 4 | 1.099 | 1.486 | 0.210 | 0.044 |
| Residual | 94.682 | 128 | 0.740 | | | |

*Note.* Type III Sum of Squares

**Table S12.** Descriptive Statistics (ANOVA) - SCL

| Condition | Time | Mean | SD | N |
|---|---|---|---|---|
| Threat | 1 | 0.920 | 0.972 | 33 |
| | 2 | 0.548 | 0.718 | 33 |
| | 3 | 0.256 | 0.723 | 33 |
| | 4 | -0.200 | 0.792 | 33 |
| | 5 | -0.042 | 1.038 | 33 |
| Safe | 1 | -5.102e -4 | 0.974 | 33 |
| | 2 | -0.187 | 0.790 | 33 |
| | 3 | -0.307 | 0.901 | 33 |
| | 4 | -0.429 | 0.838 | 33 |
| | 5 | -0.558 | 0.754 | 33 |

**Table S13. Unilateral** Paired Samples T-Test (Condition) for SCL

| | | | | | | 95% CI for Mean Difference | | | 95% CI for Cohen's d | |
|---|---|---|---|---|---|---|---|---|---|---|
| Condition | t | df | p | Mean Difference | SE Difference | Lower | Upper | Cohen's d | Lower | Upper |
| Threat - Safe | 5.942 | 32 | < .001 | 0.593 | 0.100 | 0.424 | ∞ | 1.034 | 0.672 | ∞ |

*Note.* Student's t-test.

*Note.* All tests, hypothesis is measurement one greater than measurement two.

**Table S14.** Descriptive Statistics (T-Test) - SCL

| Condition | N | Mean | SD | SE |
|---|---|---|---|---|
| Threat | 33 | 0.296 | 0.286 | 0.050 |
| Safe | 33 | -0.296 | 0.286 | 0.050 |

***Subjective Anxiety***

Repeated Measures ANOVA on Subjective Anxiety previously described on the main manuscript revealed an interaction between Condition and Time. This interaction is graphically represented in Figure S1.

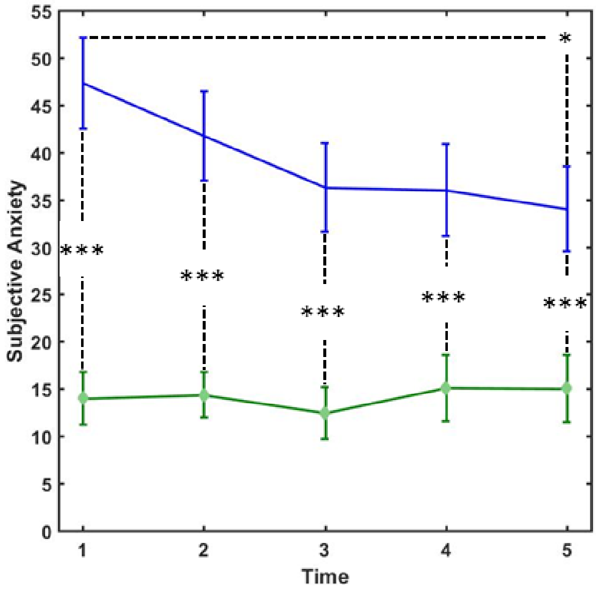


**Figure S1:** Mean of Subjective anxiety during each Threat and Safe blocks (+/- SEM). *** = p < 0.001; ** = p < 0.01; * = p < 0.05; n.s. = p > 0.05.

**Table S15.** Within Subjects Effects of Repeated Measures ANOVA (Condition & Time) on Subjective Anxiety

| | Sphericity Correction | Sum of Squares | df | Mean Square | F | p | η² |
|---|---|---|---|---|---|---|---|
| Condition | None | 51113 | 1.000 | 51113.5 | 47.844 | < .001 | 0.599 |
| | Greenhouse-Geisser | 51113 | 1.000 | 51113.5 | 47.844 | < .001 | 0.599 |
| Residual | None | 34187 | 32.000 | 1068.3 | | | |
| | Greenhouse-Geisser | 34187 | 32.000 | 1068.3 | | | |
| Time | None | 1901 [a] | 4.000 [a] | 475.1 [a] | 2.472 [a] | 0.048 [a] | 0.072 |
| | Greenhouse-Geisser | 1901 [a] | 2.324 [a] | 817.7 [a] | 2.472 [a] | 0.083 [a] | 0.072 |
| Residual | None | 24603 | 128.000 | 192.2 | | | |
| | Greenhouse-Geisser | 24603 | 74.377 | 330.8 | | | |
| Condition ✻ Time | None | 2117 | 4.000 | 529.2 | 4.046 | 0.004 | 0.112 |
| | Greenhouse-Geisser | 2117 | 3.311 | 639.2 | 4.046 | 0.007 | 0.112 |
| Residual | None | 16739 | 128.000 | 130.8 | | | |
| | Greenhouse-Geisser | 16739 | 105.958 | 158.0 | | | |

*Note.* Type III Sum of Squares

[a] Mauchly's test of sphericity indicates that the assumption of sphericity is violated (p < .05).

**Table S16.** Descriptive Statistics (ANOVA) – Subjective Anxiety

| Condition | Time | Mean | SD | N |
|---|---|---|---|---|
| Threat | 1 | 47.30 | 26.79 | 33 |
| | 2 | 41.73 | 25.79 | 33 |
| | 3 | 36.30 | 26.05 | 33 |
| | 4 | 36.03 | 26.69 | 33 |
| | 5 | 34.03 | 25.58 | 33 |
| Safe | 1 | 14.00 | 15.38 | 33 |
| | 2 | 14.36 | 13.21 | 33 |
| | 3 | 12.45 | 14.99 | 33 |
| | 4 | 15.09 | 19.11 | 33 |
| | 5 | 15.03 | 19.40 | 33 |

**Table S17.** Unilateral Paired Samples T-Test (Condition) for Subjective Anxiety

| Condition | | t | df | p | Mean Difference | SE Difference | 95% CI for Mean Difference | | Cohen's d | 95% CI for Cohen's d | |
|---|---|---|---|---|---|---|---|---|---|---|---|
| | | | | | | | Lower | Upper | | Lower | Upper |
| Threat | - Safe | 6.917 | 32 | < .001 | 24.89 | 3.599 | 18.80 | ∞ | 1.204 | 0.819 | ∞ |

*Note.* Student's t-test.

*Note.* All tests, hypothesis is measurement one greater than measurement two.

**Table S18.** Descriptive Statistics (T-Test) – Subjective Anxiety

| Condition | N | Mean | SD | SE |
|---|---|---|---|---|
| Threat | 33 | 39.08 | 23.21 | 4.041 |
| Safe | 33 | 14.19 | 12.80 | 2.229 |

**Table S19.** Post Hoc Comparisons - Subjective Anxiety during Threat Blocks

| | | Mean Difference | SE | t | $p_{bonf}$ |
|---|---|---|---|---|---|
| 1 | 2 | 5.576 | 2.889 | 1.930 | 0.625 |
| | 3 | 11.000 | 4.082 | 2.695 | 0.111 |
| | 4 | 11.273 | 3.758 | 2.999 | 0.052 |
| | 5 | 13.273 | 3.892 | 3.411 | 0.018 |
| 2 | 3 | 5.424 | 2.421 | 2.240 | 0.321 |
| | 4 | 5.697 | 2.351 | 2.424 | 0.212 |
| | 5 | 7.697 | 3.135 | 2.455 | 0.197 |
| 3 | 4 | 0.273 | 3.259 | 0.084 | 1.000 |
| | 5 | 2.273 | 4.176 | 0.544 | 1.000 |
| 4 | 5 | 2.000 | 2.768 | 0.722 | 1.000 |

**Table S20.** Post Hoc Comparisons - Subjective Anxiety during Safe Blocks

| | | Mean Difference | SE | t | $p_{bonf}$ |
|---|---|---|---|---|---|
| 1 | 2 | -0.364 | 2.711 | -0.134 | 1.000 |
| | 3 | 1.545 | 2.663 | 0.580 | 1.000 |
| | 4 | -1.091 | 3.754 | -0.291 | 1.000 |
| | 5 | -1.030 | 3.884 | -0.265 | 1.000 |
| 2 | 3 | 1.909 | 2.377 | 0.803 | 1.000 |
| | 4 | -0.727 | 3.053 | -0.238 | 1.000 |
| | 5 | -0.667 | 3.195 | -0.209 | 1.000 |
| 3 | 4 | -2.636 | 2.590 | -1.018 | 1.000 |
| | 5 | -2.576 | 2.311 | -1.115 | 1.000 |
| 4 | 5 | 0.061 | 1.920 | 0.032 | 1.000 |

**Table S21.** One Sample T-Test Subjective Anxiety (Threat minus Safe) across Time

| | | | | | 95% CI for Mean Difference | | | 95% CI for Cohen's d | |
|---|---|---|---|---|---|---|---|---|---|
| Time | t | df | p | Mean Difference | Lower | Upper | Cohen's d | Lower | Upper |
| 1 | 7.090 | 32 | < .001 | 33.30 | 25.35 | ∞ | 1.234 | 0.845 | ∞ |
| 2 | 5.924 | 32 | < .001 | 27.36 | 19.54 | ∞ | 1.031 | 0.669 | ∞ |
| 3 | 5.994 | 32 | < .001 | 23.85 | 17.11 | ∞ | 1.043 | 0.680 | ∞ |
| 4 | 4.779 | 32 | < .001 | 20.94 | 13.52 | ∞ | 0.832 | 0.493 | ∞ |
| 5 | 4.476 | 32 | < .001 | 19.00 | 11.81 | ∞ | 0.779 | 0.446 | ∞ |

*Note.* Student's t-test.

##### *Intra-individual Correlation*

**Table S22.** One Sample T-Test for Intra-individual Correlation (r to z Fischer)

| | | | | | 95% CI for Mean Difference | | | 95% CI for Cohen's d | |
|---|---|---|---|---|---|---|---|---|---|
| | t | df | p | Mean Difference | Lower | Upper | Cohen's d | Lower | Upper |
| Correlation | 4.703 | 32 | < .001 | 0.354 | 0.201 | 0.508 | 0.819 | 0.419 | 1.209 |

*Note.* Student's t-test.

**Table S23.** Descriptive Statistics (T-Test) – Intra-individual Correlation

| | N | Mean | SD | SE |
|---|---|---|---|---|
| Correlation | 33.00 | 0.354 | 0.433 | 0.075 |
